# Peptide Terminus Tilting: an Unusual conformational transition of MHC Class I Revealed by Molecular Dynamics Simulations

**DOI:** 10.1101/219873

**Authors:** Yeping Sun, Po Tian

## Abstract

A conventional picture for major histocompatibility complex class I (MHCI) antigen presentation is that the terminal anchor residues of the antigenic peptide bind to the pockets at the bottom of the MHC cleft, leaving the central peptide residues exposed for T cell antigen receptor (TCR) recognition. However, in the present study, we show that in canonical or accelerated molecular dynamics (MD) simulations, the peptide terminus in some immunodominant peptide-MHCI (pMHCI) complexes can detach from their binding pockets and stretch outside the MHC cleft. These pMHCI complexes include the complex of the H-2K^b^ and the lymphocytic choriomeningitis virus (LCMV) gp33 peptide, and the complex of the HLA-A*0201 and the influenza A virus M1 peptide. The detached peptide terminus becomes the most prominent spot at the pMHC interface, and so can serves as a novel TCR recognition target. Thus, peptide terminus detaching may be a novel mechanism for MHC antigen presentation.

## INTRODUCTION

Since Bjorkman et al. reported the crystal structures of HLA-A2 in 1987 (Bjorkman et al., 1987a, b), accumulating crystallographic data of peptide-MHC class I (pMHCI) and pMHCI-TCR complexes have defined this general principal of MHC antigen presentation: the MHC usually binds peptides of 8–10 residues in length with an extended conformation whose termini are buried in specific pockets that differ from allele to allele. This binding mode often makes the peptide central region protrude out of the MHC cleft and into the solvent (Gras et al., 2012; Rudolph et al., 2006). T cell receptors (TCRs) dock diagonally on pMHCIs with the extremely variable CDR3s generally located over the center of the pMHCI surface and make contacts with the central peptide residues as well as with the MHC α helices, while relatively conserved CDR1s and CDR2s contacts MHC α helices (Mazza and Malissen, 2007; Scott-Browne et al., 2011). Although in a few cases MHCI can bind long peptides up to thirteen amino acids, these long peptides are also anchored to the MHCI cleft by their N- and C-termini, and their central regions are forced to adopt bugled conformations (Speir et al., 2001). This basic paradigm has not been seriously challenged in spite of all the minor conformational variations reported.

However, crystal structures usually provide very limited information on molecular dynamics (MD). Internal dynamics and structural flexibility are central features of many biological processes; even small amount of structural flexibility can profoundly influence molecular recognition, signaling and, ultimately, biological function (Dror et al., 2012; Shaw et al., 2010; Smock and Gierasch, 2009), and this is also true for the recognition between the pMHCI and the TCR (Kass et al., 2014; Knapp et al., 2015; Morikis and Lambris, 2004). MD simulation is one of the most effective tools to study MD of biological molecules. The first large scale MD simulation of pMHCI was done on HLA-A*0201 bound to a decameric tumor-specific antigenic peptide GVYDGREHTV, which revealed that the peptide was less tightly bound in the peptide binding domain than the entire MHC molecule (Wan et al., 2004). Many of the following MD simulations of pMHC focused on the correlation between MHC polymorphism and pMHC dynamics. For example, simulations of an 13-mer peptide (LPEPLPQGQLTAY) bound to MHC class I molecules HLA-B*3501 and HLA-B*3508 showed that the 13-mer peptide in the SB27 TCR-reactive complex (HLA-B*3508–13-mer) exhibited lower flexibility compared to the peptide in the non-active HLA-B*3501–13-mer complex (Reboul et al., 2012). Although B*3501 and B*3508 have only one different residue (R156 for B*3508 and L156 for B*3501) and crystal of these two pMHC complexes are almost identical, B27 TCR can discriminate them (Tynan et al., 2005a; Tynan et al., 2005b). Thus, different dynamics of these two pMHC provided persuasive explanation for their TCR reactivity.

Regulation of the dynamics of the MHC molecule by the bound peptide has also been reported. One example is that variations in the MART-1 melanoma peptide (AAGIGILTV→ALGIGILTV) induced flexibility of the α1 and α2 helices of HLA-A*0201, and thus weakened TCR ligation, and therefore, immunogenicity (Insaidoo et al., 2011). In our own study, we showed that HLA-A*0201 in complex with two avian influenza virus H5-specific CTL epitope, RI-10 (RLYQNPTTYI) and KI-10 (KLYQNPTTYI), differed significantly in their dynamics. KI-10-HLA-A*0201 complex processed much higher flexibility than RI-10-HLA-A*0201, and the higher dynamics of KI-10-HLA-A*0201 was associated with its lower antigenicity (Sun and Liu, 2015).

In the above studies, higher structural fluctuation or flexibility at the TCR recognition interface of pMHC, including the peptide and MHC α helices seem to frustrate TCR ligation. This seems to be coherent with the findings in the instance of antigen-antibody recognition (Singharoy et al., 2013). Paradoxically, flexibility at the pMHC-TCR interaction interface, especially at CDR loops, and also at the peptide and the MHC, plays a prominent role in binding (Borbulevych et al., 2011; Borbulevych et al., 2009). The unligated Tel1p (MLWGYLQYV) and Tax (LLFGYPVYV) in complex with HLA-A*0201 were sequence and structural mimics, but the interface they formed with A6 TCR substantially different, with conformational change in the peptide, the TCR, and the HLA-A*0201 α2 helix. The differences were attributable to peptide and MHC molecular motion present in Tel1p-HLA-A*0201 but absent in Tax-HLA-A*0201, which was revealed by MD simulations (Borbulevych et al., 2009).

Although numerous MD simulation studies have been done with pMHCI, few reported large-scale conformational changes for these molecules. In the MD simulations reported by most of these studies, the basic binding pattern between the peptide and the MHC molecule remains the same: the anchor residues at both termini of the peptide kept buried in the binding pockets at the bottom of antigen binding cleft of the pMHC molecules. However, in our MD simulations with the pMHCI molecules in which the antigenic peptides are immunodominant viral epitopes, one terminus of the peptides in some of these molecules can detach from the MHC binding pockets and float toward outside of the MHCI groove. This phenomenon may represent an unusual antigen presentation strategy of MHC molecules.

## Results

**Peptide N-terminus detaching in the complex of the H-2K^b^ and the lymphocytic choriomeningitis virus (LCMV) gp33 (33-41) peptide (H-2K^b^-gp33) in canonical MD (cMD) simulations**

### The gp33 octapeptide

The LCMV-infected mouse is a classic model for cell-mediated immunity characterized by a CD8^+^ T cell response in which the majority of the CTLs are directed against a few immunodominant epitopes in viral proteins. One of these epitopes is gp33 peptide (KAVYNFATM) in the context of H-2K^b^ alleles. The structure of H-2K^b^-gp33 is very unusual. Unlike the vast majority of peptides known to bind to MHCI molecules through a network of hydrogen bond interactions between their amino and carboxyl termini and the conserved amino acids in the A- and F-pockets at the ends of the peptide binding cleft of the MHCI, only eight residues of the nonameric gp33 epitope (residues p2A to p9M) occupy the peptide binding groove of H-2K^b^, leaving p1K at the N terminus outside the cleft (Achour et al., 2002; Velloso et al., 2004).

We carried out a 100 ns MD simulation on the H-2K^b^-gp33 complex. We first used the H-2K^b^-gp33 crystal structural model (PDB code: 1n59) as the initial structure for the simulation. The p1K residue of the peptide is missing in the model because its electron density is invisible in the crystal structure (Figures 1A and 1B). The root mean square deviation (RMSD) values of the whole protein became roughly converged after 40 ns in the 100 ns simulation (Figure 1C). In the root mean square fluctuation (RMSF) plot for each residue in the pMHCI heavy chain (HC) and the peptide, the N-terminal residues of the peptide, p2A and p3V, have the highest RMSF values, suggesting that they were the most fluctuating residues among all the residues shown here (Figure 1D). Intriguingly, the fluctuation of these two residues can be intuitively reflected by the motions in the simulation trajectory. They detached from the H-2K^b^ bottom and moved upwards a few nanoseconds after initiation of the simulation, and then fully stretched out of the peptide binding cleft of the H-2K^b^ molecule and flapped around (Movie S1).

**Figure 1.**
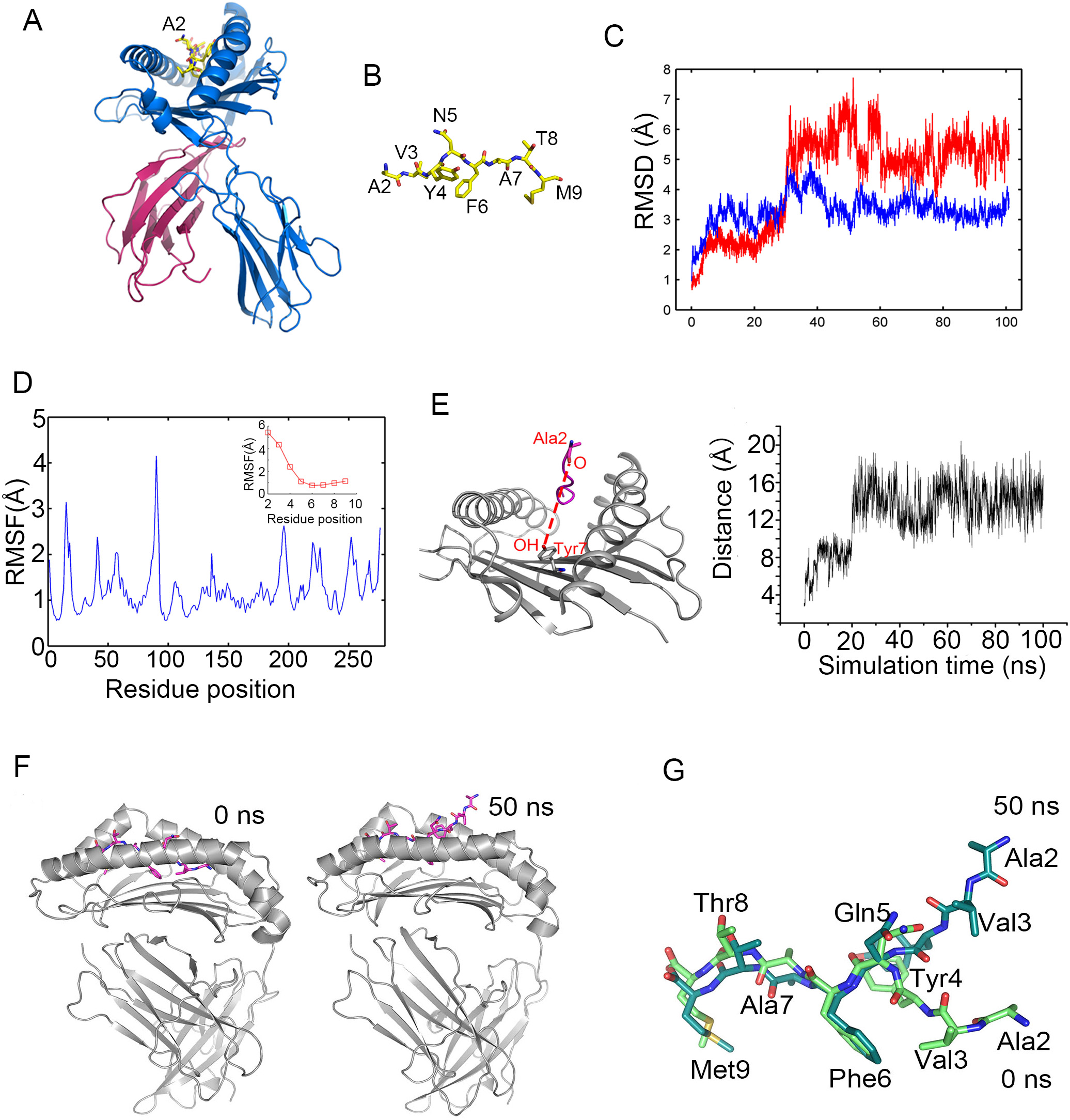
N-terminus detaching in the H2-K^b^/gp33 (8-mer) in cMD simulation. (A) The crystal structure of H2-K^b^/gp33 complex (PDB ID: 1N59). (B) The conformation of the gp33 peptide in the crystal structure of the H2-K^b^/gp33 complex. (C) RMSD of the whole H2-K^b^/gp33 protein and the gp33 octapeptide. (D) RMSF of each residue in the H2-K^b^/gp33 HC and the peptide. (E) Distance between the O atom in Ala2 of gp33 and the OH atom in Tyr7 of H2-K^b^ molecule. (F) Comparison of the H2-K^b^/gp33 structures at 0 and 50 ns. (G) Comparison of the peptide conformations in aligned H2-K^b^/gp33 structures at 0 and 50 ns.

The upward movement of the peptide N-terminus can be measured by distances between the O atom of Ala2 within the gp33 peptide and the OH atom of Tyr7 within the H-2K^b^ HC, which is located in the Ala2 binding pocket (pocket A) (Figure 1E, the left panel). At 0 ns, the two entities were close to each other, with a distance of3.15 Å. Shortly after the beginning of the simulation, however, the distance between them increased to approximately 8 Å and fluctuated around this level until 20 ns, when the distance suddenly further jumped to 15 Å and then fluctuated at this level for the remaining simulation time (Figure 1E, the right panel). Here we just illustrate the peptide N-terminus conformational change by comparing the structures snapshots at the beginning of the simulation (0 ns) and 50 ns. At 0 ns, both the N- and C- termini bound to their respective binding pockets and buried in the peptide binding cleft, but at 50 ns, the peptide N-terminal p2A and p3V were totally exposed outside the peptide binding cleft (Figure 1F). Superposition of the two snapshots shows that the postures of both the main chains and the side chains of the six C-terminal residues (Tyr4 to Met 9) of the peptide at these two time points are very similar, while those of the two residues at the N- terminus, p2A and p3V, are dramatically different. At 50 ns the main chain of these two residues bends upwards and leaves far away from their initial position at 0 ns (Figure 1G).

### The gp33 nonapeptide

To examine whether the N-terminus detaching would occur in the presence of the first residue, Lys1 of the gp33 peptide, we modeled the structure of the gp33 nonapeptide in complex with the H2-K^b^ by Chimera program (Pettersen et al., 2004) and performed MD simulations under the same conditions with the gp33 octapeptide in complex with the H2-K^b^, as described above. After the system was subjected to energy minimization, 1 ns C_α_ constraint simulation at 310 K and 1 ns equilibrium simulation without constraints, we obtained the 0 ns protein structure (Figure 2A). In this structure, residues 2 to 9 of the nonapeptide overlap well with the octapeptide in the crystal structure, and the modeled p1K points towards upwards without occupying any classic peptide binding pocket in the cleft bottom. Just like in the simulation of the H2-K^b^-gp33 octapeptide, the RMSD of the H2-K^b^-gp33 nonapeptide complex become converged after 40 ns (Figure 2B). And the RMSF values of peptide residues p1K and p2A are the highest among all the residues shown (Figure 2C). As we expected, we do observe peptide N-terminus detaching in this modeled complex (movie S2). Again, we measured the distance between the O atom of peptide Ala2 and the OH atom of H-2K^b^ HC Tyr7 as an index of peptide N-terminus detaching and upwards movement. From 0 to 5 ns this distance was about 5 Å, then it decreased slightly to around 4 Å and kept at this level until 28 ns, after which the distance rapidly increased to and fluctuated around 12 Å until the end of the simulation (Figure 2D). The peptide RMSD-based clustering analysis shows that the peptide conformations in the 100 ns MD trajectory are grouped into two clusters. Cluster 1 accounts for 24.3 % of the trajectory and Cluster 2 occupies 51.2 % of that. Cluster 1 occurs before 28 ns and Cluster 2 appeared after that time point, which suggested the time point 28 ns is a watershed of the conformational space. (Figure 3A). Comparing the representative structures of these two clusters shows the sharp contrast of the peptide N-termini conformations (Figures 3B and 3C). Just like in the structure model at the 0 ns, the p2A and p3V in the representative structure of Cluster 1 bind to the bottom of H-2K^b^ cleft, but in the representative structure of Cluster 2, the p2A and p3V detach from their original binding sites and move upwards, making the p1K more exposed to the solvent. By contrast, the remaining residues (p4Y to p9M) of the peptide superpose well in these two representative structures (Figure 3C).

**Figure 2.**
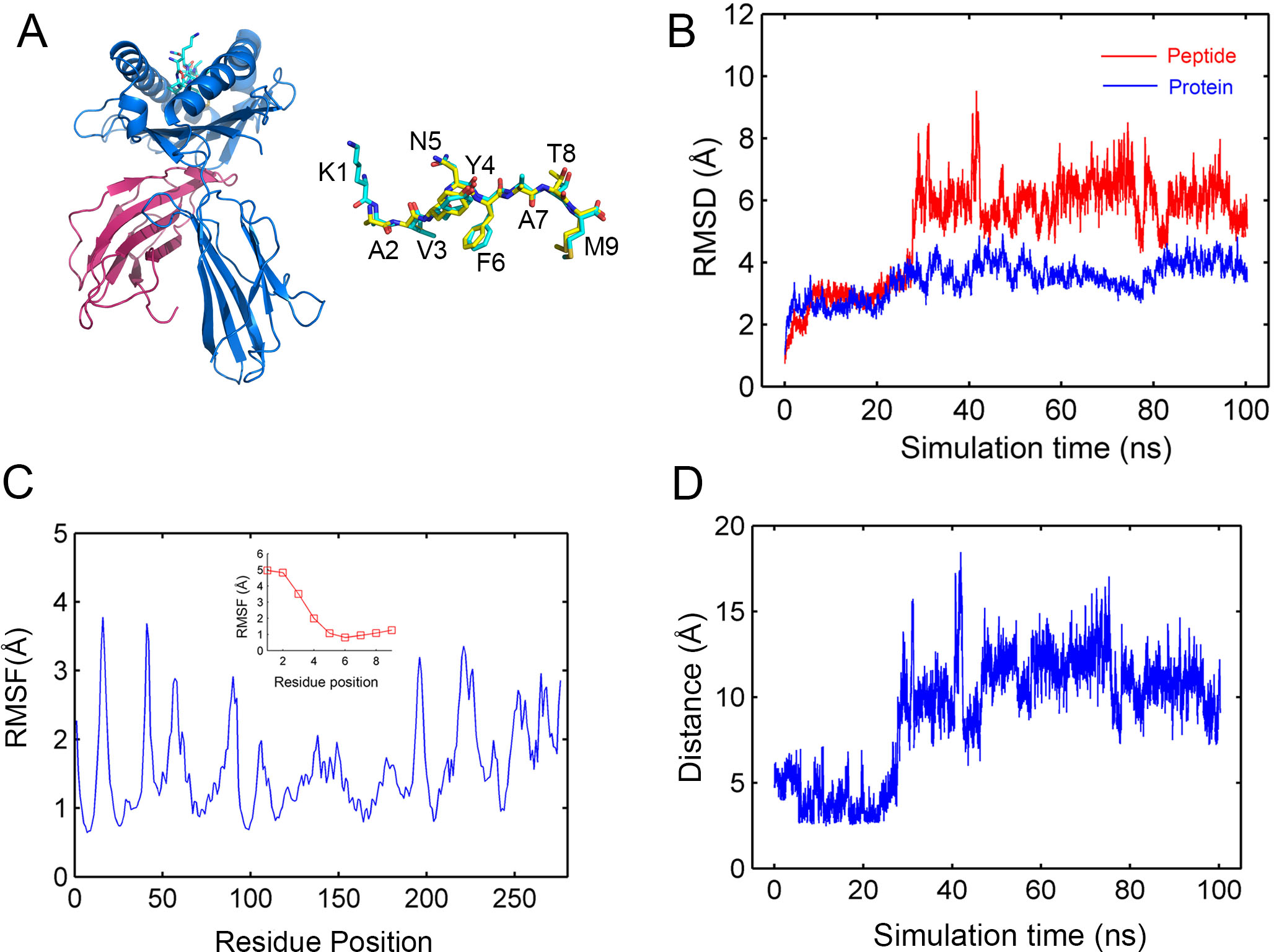
N-terminus detaching in the H2-K^b^/gp33 (9-mer) in cMD simulation. (A) The overall structure of the modeled H2-K^b^/gp33 (9-mer) (left panel) and the aligned 9-mer and 8 mer gp33 peptide from the modeled H2-K^b^/gp33 (9-mer) and the crystal structure of H2-K^b^/gp33 (8-mer). (B) RMSD of the whole H2-K^b^/gp33 protein and the gp33 nonapeptide. (C) RMSF of each residue in the H2-K^b^/gp33 HC and the peptide. (D) Distance between the O atom in Ala2 of gp33 and the OH atom in Tyr7 of H2-K^b^ molecule.

**Figure 3.**
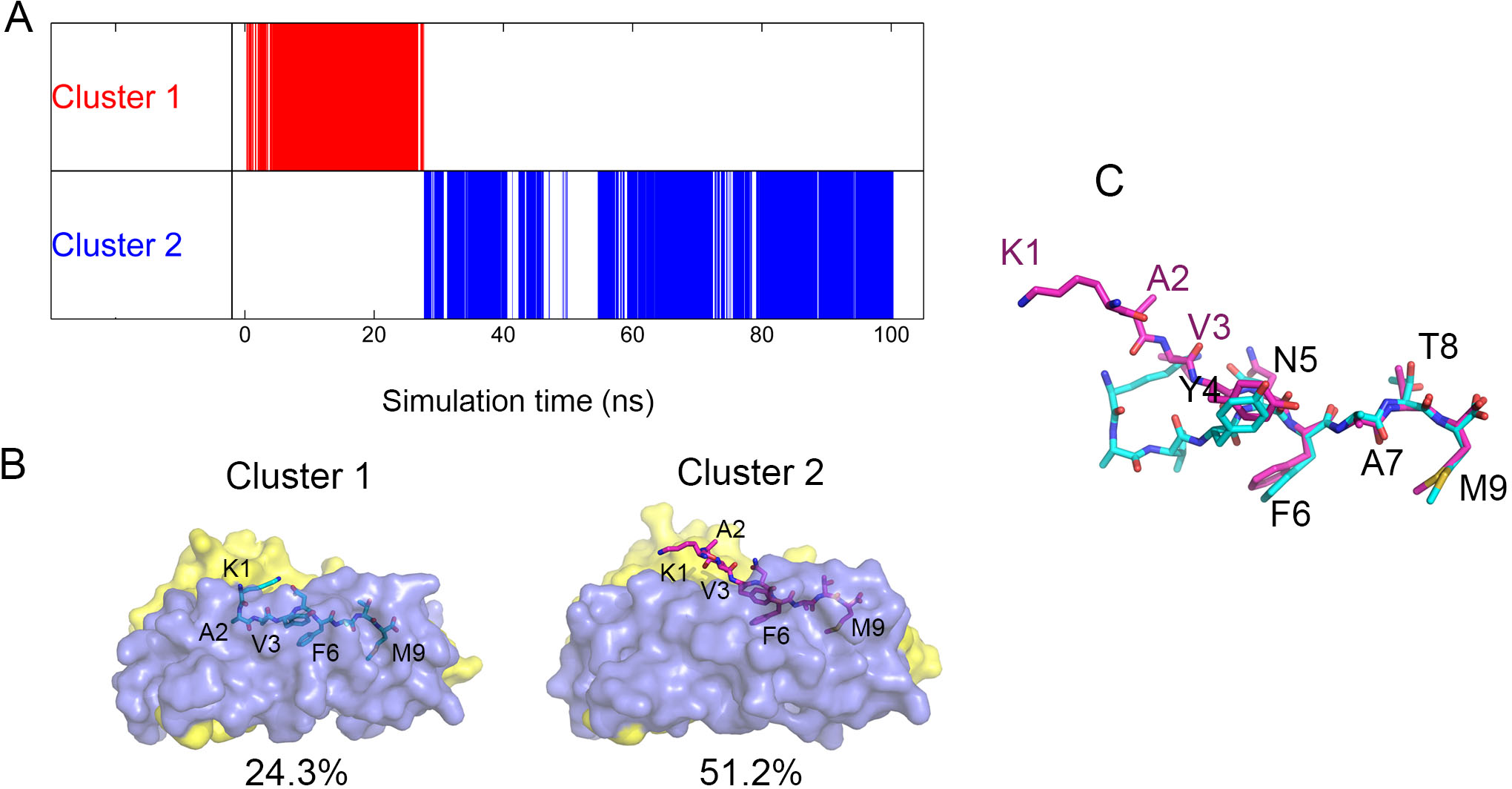
Cluster analysis of the peptide conformations in the H2-K^b^/gp33 (9-mer) in the cMD simulation. (A) The snapshot distribution of the two peptide conformation clusters, Cluster 1 and 2, during the 100 ns simulation time. (B) The representative structure of Cluster 1 and 2. Only the peptide, the α1 and α2 domains of the H2-K^b^ are shown. The α1 and α2 domains are indicated as cyan and yellow, respectively. (C) The aligned peptides in Cluster 1 and 2.

Next, we attempt to examine the underlying mechanisms that lead to peptide N-terminus detaching in this pMHC complex. By calculating the interaction energy (Figure 4A) between every residue of the peptide and the H2-K^b^ HC in the representative structure of Cluster 1 we know that the interaction energy between p1K and the H2-K^b^ HC is highly positive, and those between p2A and p3V and H2-K^b^ HC are small negative values. In contrast, the interaction energy between two C-terminal residues and the H2-K^b^ HC is highly negative. These data suggest that the binding between the peptide N-terminus is energetically much less favorable than the peptide C-terminus and so provide clues to understand why N-terminus rather than C-terminus of the peptide tilted in the MD simulation. They also suggest that the peptide Tyr4, the secondary anchor residue whose interaction energy is negatively much higher than that of the p2A and p3V, also plays an important role in holding the peptide into the H2-K^b^ HC cleft, as the peptide C-terminus does.

**Figure 4.**
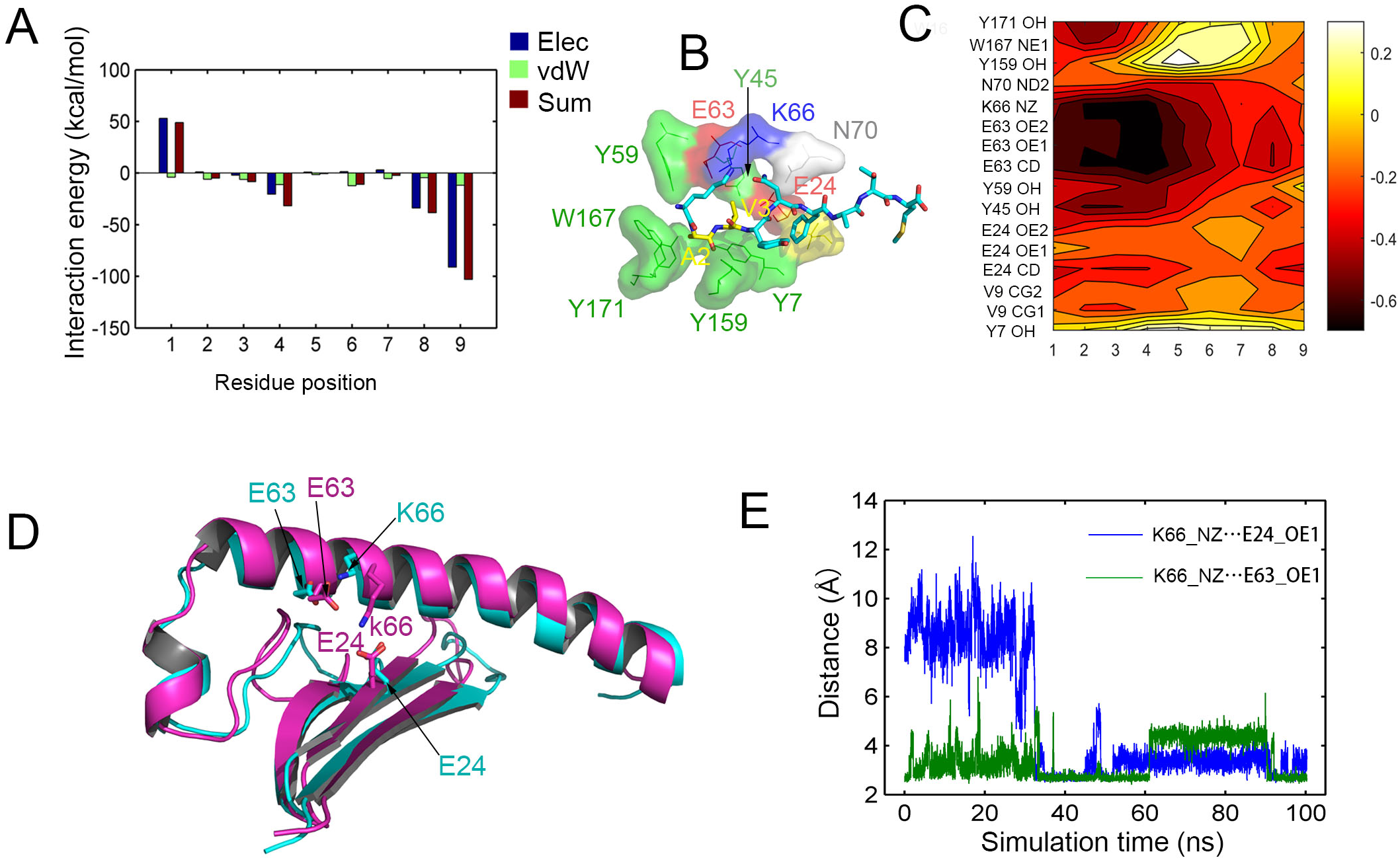
The interactions between the nonapeptidic gp33 and the H2-K^b^ in the cMD simulation. (A) The interaction energy between each individual residue of the peptide and the H2-K^b^. (B) The H2-K^b^ residues that contact the N-terminal four residues in the gp33 peptide. The H2-K^b^ residues are shown as surface and the peptide residues are shown as sticks. The aromatic, acidic, basic and aliphatic residues in the H2-K^b^ are indicated with green, red, blue and yellow, respectively. The aliphatic Ala2 and Val3 in the gp33 peptide are also indicated with yellow. (C) The cross-relation matrix of the H2-K^b^ atoms and the peptide Cα-atom. (D) Alignment of the representative structures of Cluster 1 (cyan) and 2 (purple) showing the change of relative positions of H2-K^b^ E24 and K66. In Cluster 1 the two residues are far from each other, but in Cluster 2 they come closer to each other and from a salt bridge. (E) The distance H2-K^b^ K66 N atom and E24 OE1 atom, and the distance between H2-K^b^ K66 N atom and E63 OE1 atom.

In the representative structure of Cluster 1, the peptide residue p2A is surrounded by five aromatic residues Y7, Y59, Y159, W167, Y171, as well as an acidic residue, E63 from H2-K^b^ HC; and the side chain of the peptide residue p3V points to the B-pocket formed by H2-K^b^ HC residues Y7, V9, E24, Y45, N70 (Figure 4B). The peptide N-terminus detaching occurs between 24 to 34 ns in the MD trajectory (figure 3D), so we calculated the dynamics cross-correlation matrix (DCCM) between these residues of the H2-K^b^ HC and the individual peptide residues using this segment of the trajectory to examine whether there is any movement correlation between the peptide N-terminal residues and their interacting residues in H2-K^b^ HC. For the peptide residues we used their CA atoms, but unlike the peptide N-terminal residues, CA atoms of those H2-K^b^ residues just vibrate in the backbone without large scale of movement, so we use the terminal atoms of their side chain for the analysis. The results show that the side chains of H2-K^b^ HC K66 and E63 have high negative correlation with the five N-terminal residues of the peptide (Figure 4). By observing the 20 to 44 ns of the simulation trajectory carefully we can see that when p2A and p3V began to detach and move upwards at about 22 ns, the side chain H2-K^b^ HC K66 stretches downwards simultaneously and forms a salt bridge with E24 at the bottom of the B-pocket and occupy the original position of p3V (Movie S3). This makes us hypothesize that the trend of formation of the salt bridge between K66 and E24 is a driving force of peptide N-terminal detach. Superposition of the representative structures of Cluster 1 and 2 of the simulation trajectory shows that there is an obvious displacement for the side chain of K66. In the representative structures of Cluster 1 the side chain of K66 is far from the side chain of E24, but in the representative structures of Cluster 2, it forms a salt bridge with E24 (Figure 4D). At the beginning of the simulation trajectory, the distance between K66 NZ atom and the E24 OE1 atom was around 8 Å, but it drops to 2.6 Å at 32 ns, which suggest the formation of the salt bridge, and kept being less than 4 Å until the end of the simulation except a transient shooting up at around 48 ns.

### Peptide terminus detaching in some other immunodominant pMHC complexes in accelerated MD (aMD) simulations

To explore whether the peptide terminus detaching phenomenon is common in other pMHCI complex molecules, we performed 40-100 ns cMD simulation in several other representative pMHC complexes. These pMHCI molecules, include influenza A virus M1 (58-66) in complex with HLA-A*0201 (HLA-A2-M1) (PDB ID: 1HHI), Human immunodeficiency virus-1 (HIV-1) p24 Gag (263-272) in complex with HLA-B*2705 (HLA-B27-Gag) (PDB ID: 4G9D), HIV-1 Nef (134-143) in complex with HLA-A*2402 (HLA-A24-Nef) (PDB ID: 3VXN), and influenza A virus NP (418-426) in complex with HLA-B*3501 (HLA-B35-NP) (PDB ID: 3LKN), are all composed of immunodominant epitopes from different viruses and different HLA alleles. We didn’t observe obvious peptide terminus detaching among them in our cMD simulations (Figure 5).

**Figure 5.**
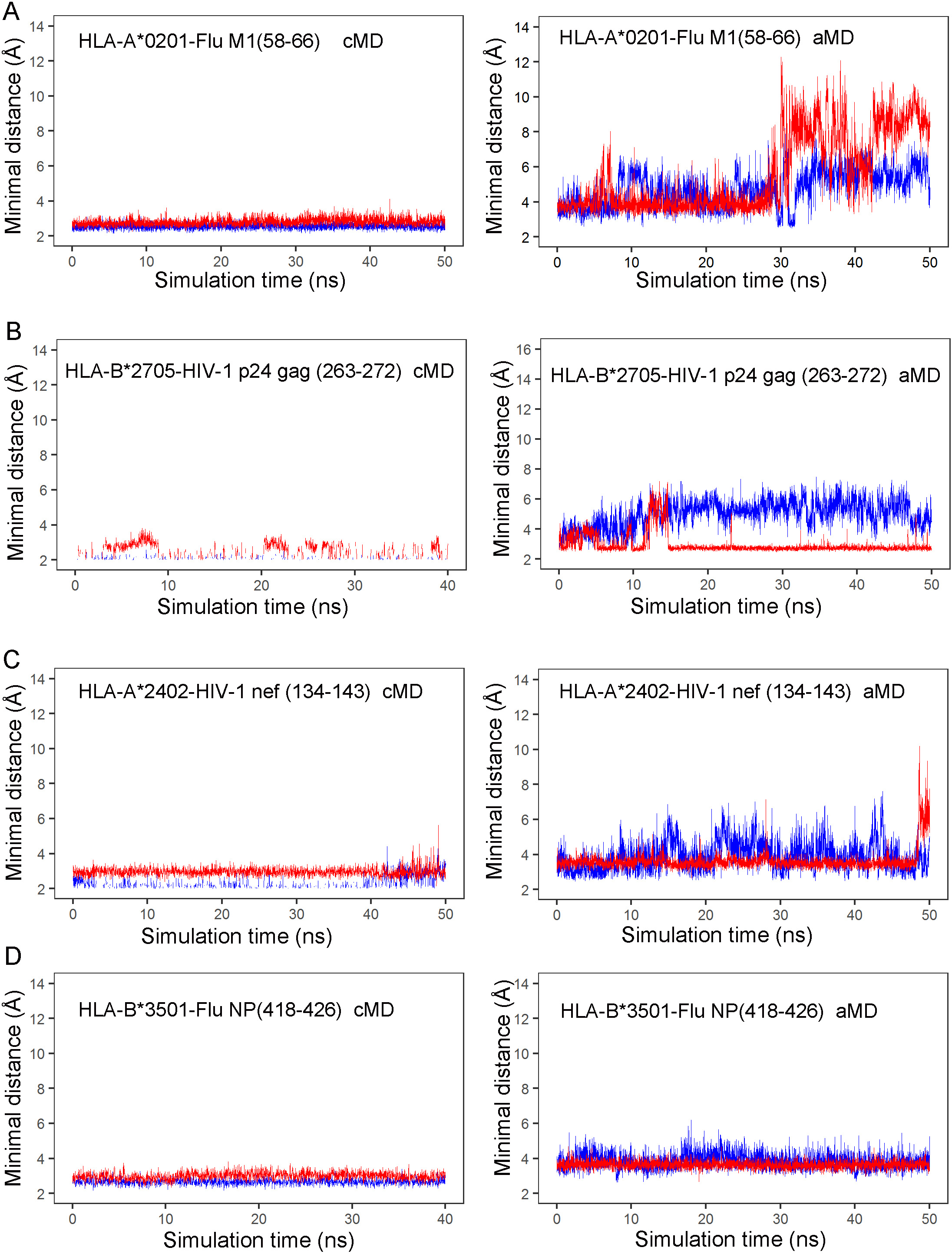
Peptide terminus detaching in cMD (the left panel) and aMD (the right panel) simulations with several immunodominant pMHCIs in viral infection. (A) influenza A virus M1 (58-66) in complex with HLA-A*0201 (PDB ID: 1HHI). (B) Human immunodeficiency virus-1 (HIV-1) p24 Gag (263-272) in complex with HLA-B*2705 (PDB ID: 4G9D). (C) HIV-1 Nef (134-143) in complex with HLA-A*2402 (PDB ID: 3VXN). (D) influenza A virus NP (418-426) in complex with HLA-B*3501 (PDB ID: 3LKN). The blue and red line represent the minimal distance from the Cα atoms of the first and the last peptide residues to the bottom of the MHCI molecule, respectively.

A drawback of cMD simulation is that the system may be trapped in a potential basin and a conformational transition is thereof difficult to occur. So it is common that a cMD in limited simulation time scale is unsuccessful in sufficiently exploring the conformational space of biomolecules. One of the solutions to this problem is the aMD methods in which a boost energy is applied to the system. As a result, when the energy barrier is conquered, the conformational transition may appear (Hamelberg et al., 2004). Herein, we performed aMD simulations on those pMHC molecules we selected. During these simulations, extra booster energies were added onto the dihedral and the total potentials, respectively (Wang et al., 2011). We measured the minimal atom distance from the p1 residue of peptides to the bottom of the MHCI HC they bind and arbitrarily defined a minimal atom distance larger than 8 Å as the indicator of obvious peptide terminus detaching. Among these molecules we tested, we observed that in the HLA-A2-M1 complex, the C-terminus of the M1 (58-66) peptide obviously detached from its binding pocket after 30 ns. The N-terminus of this peptide also showed a trend of moving upwards, but didn’t reach a degree of obvious detaching (Figure 5A). In the HLA-B27-Gag complex, the HIV-1 Gag peptide N-terminus moved upwards, but didn’t reach the degree of obvious detaching either. For its C-terminus, only a transient upwards movement occurred at 13 ∼ 15 ns, but in most of the simulation time it bound steadily at its pocket (Figure 5B). In the HLA-A24-Nef complex, the HIV-1 Nef peptide N-terminus moved upwards but didn’t obviously detach, while its C-terminus kept steadily in its pocket until at the end of the simulation it suddenly detached (Figure 5C). In the HLA-B35-NP complex, both the N- and C-termini of the influenza A virus NP peptide bound stably to their pockets (Figure 5).

### Peptide C-terminus detaching in the HLA-A2-M1 complex

Since we observed obvious peptide C-terminus detaching in the HLA-A2-M1 complex in aMD, we next did a detailed analysis of the conformational transition in this complex. We performed a 300 ns aMD simulation using the same dual booster energy strategy as described above. During the simulation, the RMSD of the whole protein became converged after 150 ns (Figure 6A). The plot of the minimal atom distance from the peptide C-terminal residue Leu9 to the bottom of the HLA-A2 showed that the peptide C-terminus moved upwards and detached from its binding pocket and became exposed to solvent after 30 ns but rebound to its binding pocket at the end of the simulation (Figure 6B). The conformational transition of peptide C-terminus detaching and rebinding can also be observed in Movie S4. The detached peptide C-terminus can flip not only perpendicularly (in the direction of up and down), but also horizontally, so we chose a second reaction coordinate, the distance from the CA atom of the peptide Leu9 residue to the central plane (the plane passing the principal axis parallel to the MHC HC helices and the principal axis perpendicular to bottom of the MHC cleft). If the Leu9 CA atom locates just in the plane, the distance is 0. If it moves to the of the α1 helix side of the MHC HC, the distance is positive, otherwise, if it moves to the α2 helix side, the distance is negative (Figure 6C). The distance from Leu9 CA atom to the central panel is 0 at the beginning of the simulation, suggesting it located in the central plane. However, it was closer to the α2 helix between 0 and 100 ns and then move to the other side of the central plane and remained in the space between the central plane and the α1 helix until 200 ns. Between 200 ns and 300 ns, it flipped between the two helices around the central plane. We then constructed the free energy surface using the two reaction coordinates, the minimal distance from peptide Leu9 to the MHC cleft bottom and the distance from the peptide Leu9 CA atom to the central plane. The free energy surface was reweighted by Maclaurin series and is shown in Figure 6E. Two low energy regions can be identified in the free energy surface, and they correspond to the conformation of peptide C-terminus bound to its pocket and the conformation of peptide C-terminus detached from its pocket, respectively. In the representative structure for the minimal energy point in the peptide C-terminus detached conformational region, the peptide Leu9 is totally exposed at the outside of the MHC cleft and its large side chain which originally pointed downwards in the crystal structure, now turns upwards and formed an extremely prominent contact site for the TCR.

**Figure 6.**
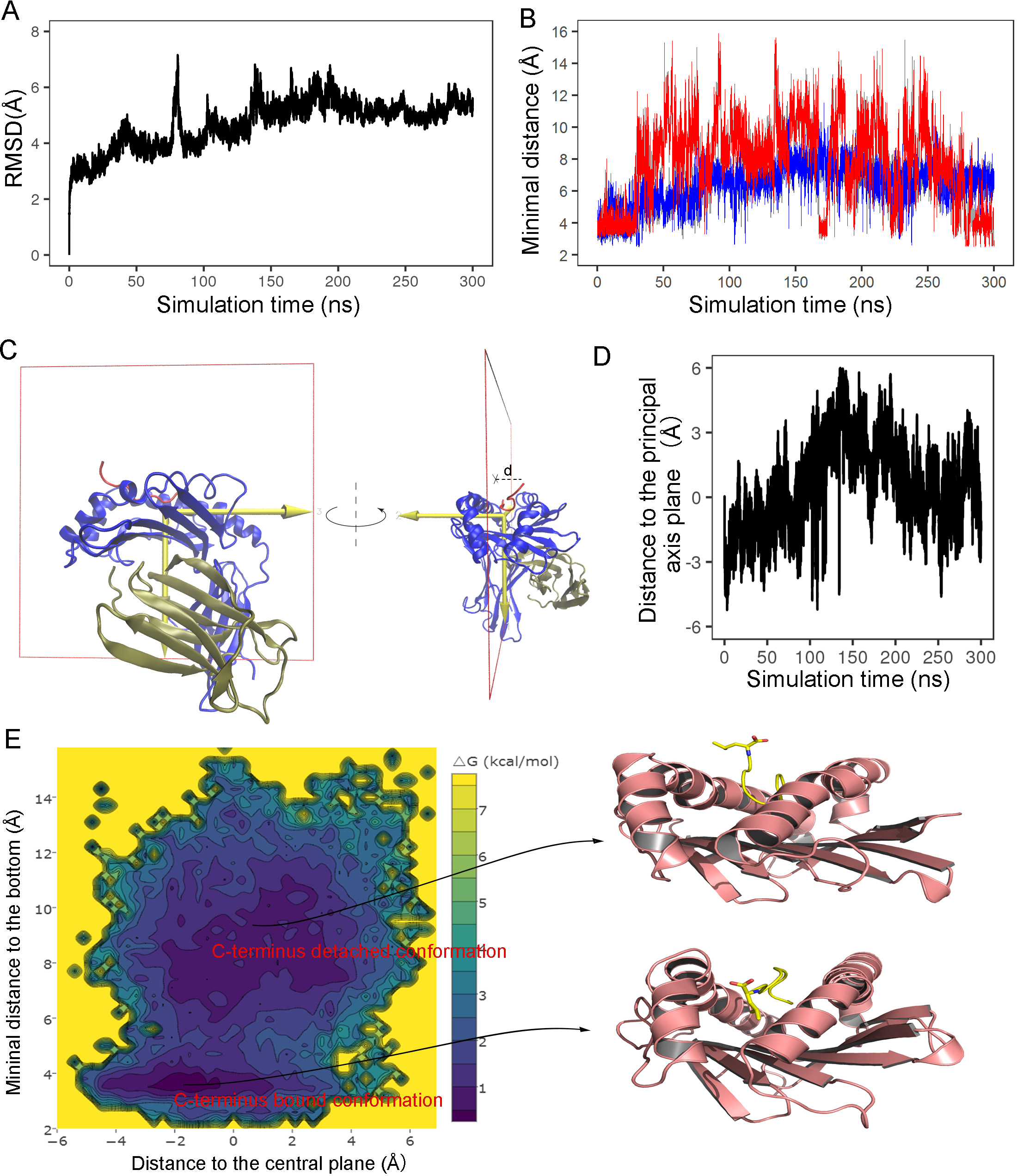
The peptide C-terminus detaching in the HLA-A2-M1 complex during the aMD simulation. (A) RMSD of the HLA-A2-M1 protein. (B) The minimal distance from the Cα atom of the first (blue) and the last (red) M1 (58-66) peptide residues to the bottom of the HLA-A2 molecule during the aMD. (C) The HLA-A2-M1 central plane (shown as the red box) which pasts the principal axis parallel to the MHC HC helices and the principal axis perpendicular to bottom of the MHC cleft (the left panel), and the distance from the CA atom of the M1 (58-66) C-terminal residue to the central plane (the right panel). (D) The distance from the CA atom of the M1 (58-66) C-terminal residue to the central plane during the aMD. (E) The unweighted free energy landscape (the left panel) and the representative structures of the two low energy regions in the free energy landscape (the right panel).

### Discussion

In the present study, we report an unusual peptide conformational transition we captured in some immunodominant pMHCI complexes in MD simulations: either the peptide N- or C-terminus can be dissociated from their binding pockets and protrude towards outside of the MHCI cleft. The peptide terminus detaching conformation can last a relatively long time during MD simulations and leave their corresponding binding pocket empty. The correct folding and stabilility of MHC is supposed to depend on optimal peptide binding, but the peptide binding to MHCI molecule without using one of its terminus has been reported. Glithero et al. reported the crystal structure of H-2D^b^ in complex with a C-terminal half (residues 5-9) of a 9-mer peptide. In this structure, the conformation of the C-terminal three peptide residues is common to its counterpart in the complex of the full-length peptide and the H-2D^b^ molecule. This crystal structure suggests that pMHC can be stable in the absence of one terminus (Glithero et al., 2006). In an extreme case, even the dipeptides can bind to the F pocket of H-2K^b^ and HLA-A*0201 molecules and stabilize them, and the dipeptides are supposed to help the MHC to fold into a peptide-receptive status that rapidly binds high-affinity peptides (Saini et al., 2013). So, it is not unexpected that peptide-terminus detaching in our simulations does not significantly affect the stability of the pMHC molecules.

The peptide terminus detaching phenomenon was not first described in the present study, but by Omasits et al. who showed that C-terminal end of an artificially constructed epitope (ARAAAAAAA) could detach from MHC and “single-site” bound to HLA-B*2505 in 1 ns MD simulations (Omasits et al., 2008). Later, the peptide terminus detaching was also mentioned for the HLA-B*2705- and HLA-B*2709- bound peptides by Narzi (Narzi et al., 2012). Unlike those studies, our work focused on virus-derived immunodominant peptides, and we gave much more detail analysis, thus making the data presented here more relevent for exploring biological significance of the peptide terminus detaching behavior.

Since the pMHCIs we studied are immunodoninant, the peptide terminus detached conformation identified in them can play a role in effective antigen presentation. The detached peptide terminus protrudes into the solvent and becomes the summit on the pMHC platform, so it can be the most approachable spot for the incoming TCR. Consequently, the pMHC may use the flexible detached peptide terminus to bind the TCR and then lead it to dock to the proper position. Based on our data, the peptide terminus detached conformation can be dominant (Figure 3B) and form low energy region in the simulation trajectories (Fig 6E), so it can play a substantive role in TCR recognition.

Among the pMHCIs we tested, only H-2K^b^ in complex with LCMV gp33 peptide displayed terminus detaching in cMD simulations, but for other pMHCIs, the peptide terminus detaching can also be revealed when proper booster energy is applied and certain energy barrier is overcome. In aMD simulations, HLA-A2-M1 the complex showed the most obvious detaching conformational transition. In the crystal structure of the HLA-A2-M1 complex, most side chains of the M1 peptide are buried in the HLA-A2 cleft and the peptide shows a featureless conformation. Consist with a previously established rule that featureless peptide selects biased TCR repertoire in contrast to featured peptide with prominently exposed side chain which induces diverse TCR repertoire (Turner et al., 2006; Turner et al., 2005), HLA-A2-M1 complex is highly prone to use a dominant public TRBV19/TRAV27(+) TCR (Valkenburg et al., 2016). However, by the next-generation-sequencing (NGS), the HLA-A2-M1-specific TCR repertoire was reveal to be broader. In addition to the dominant TRBV19, there are minor usage of other TRBV genes (such as TRBV07, 09, 28, etc.) for the TCRβ chains. The TRAV genes selected for the TCRα chains were much more diverse. In addition to TRAV27, many other TRAV genes, including TRAV12, 13, 25, 29 and 38 were commonly detected in different donors. In the crystal structure Two newly identified TCRs, LS01 and LS10, show different modes in recognition of HLA-A2-M1 (Song et al., 2017). In our aMD simulations, the C-terminus of the featureless M1 peptide in the HLA-A2 cleft can become largely exposed. This provides n potential explanation for the diverse TCR repertoire used for the HLA-A2-M1 complex.

For larger protein molecules, reweighting of an aMD simulation trajectory is difficult and there has been no successful example of recovering the canonicial ensemble from aMD simulations of a protein with nearly 300 residues like pMHCI by reweighting. We tried several different reweighting methods including exponential average, cumulant expansion and Maclaurin series (Miao et al., 2014), Only Maclaurin series gave relatively satisfying resuts wheares the reweighted free energy surfaces generated by the other two algorithms were contaminated by serious noisy. Although our simulation results remain to be verified by experiments, they still open a new window to envisage antigen presentation mechanisms pMHCI complexes.

## SUPPLEMENTARY INFORMATION

Movie S1. N-terminus tilting of H2-K^b^ bound gp33 octapeptide during the cMD simulation.

Movie S2. N-terminus tilting of H2-K^d^ bound gp33 nonapeptide during the cMD simulation.

Movie S3. The formation of the salt bridge between the H2-K^b^ E24 and K66 during the N-terminus detaching.

Movie S4. The peptide C-terminus detaching in the HLA-A2-M1 complex during the aMD simulation.

## ARTHUR CONTRIBUTIONS

Y.S. designed and performed the simulation experiments, analyzed the data and wrote the paper; P.T. supervised the research and revised the paper.

## ACKNOWLEDGMENT

This work was supported by grant from the National Natural Science Foundation of China (NSFC, Grant No. 81621091). We are grateful to Dr. Paul Chu, Professor of the Institute of Microbiology, Chinese Academy of Sciences, for his help during the preparation of the manuscript. All the MD simulations in this work were done on the Tianhe-2 supercomputer in National supercomputer center in Guangzhou, China. Both authors declare they do not have any conflict of interest.

## METHODS

### Preparation of simulation systems

Five pMHCI complex molecules are included in this study. They are H-2K^b^-gp33 (PDB ID: 1N59), HLA-A2-M1 (PDB ID: 1HHI), HLA-B27-Gag (PDB ID: 4G9D), HLA-A24-Nef (PDB ID: 3VXN), HLA-B35-NP (PDB ID: 3LKN). These structures were downloaded from Protein Database and used as the initial structures for MD simulations. If the downloaded original structures contained more than one copy of the pMHCI complex, only one copy was used. Water molecules in original structures were included in the initial structures for simulations. In the original structure for the H-2K^b^-gp33 complex, the peptide N-terminal Arginine residue is missing. This missing Arginine was modeled by Chimera program. Both the Arginine-missing and the modeled Arginine-containing version of the H-2K^b^-gp33 complex were used for MD simulations. The initial structures were solvated with water (TIP3P model) and 0.15 mM NaCl to obtain the simulation systems.

### cMD simulations

MD simulations were performed using the NAMD (Phillips et al., 2005) version 2.11 software package and the CHARMM27 force field (Foloppe and MacKerell, 2000). All chemical bonds that contain hydrogen atoms were constrained by SHAKE algorithm. For all simulations, a time step of 2 fs was used as well as periodic boundary conditions and PME for electrostatics. The simulation systems were first subjected to energy minimization, and then heated to 310 K and simulated for 1 ns with constraining Cα atoms, after which the systems were equilibrated for another 1 ns without constraints before the production runs used for analysis. Parameters used for all production simulations were: temperature 310 K, switching distance 10 Å, switching cutoff 12 Å, pair-list distance 14 Å, Langevin damping coefficient 1 ps^-1^, and Langevin pressure control with a target pressure of 1.01325 bar.

### aMD simulations

aMD simulations were performed by applying a boost potential onto the original potential of the system if the original potential is lower than a user defined threshold value, while an acceleration factor is used to optimize the modified potential curve. Herein we used the dual aMD implemented in NAMD in which two independent boost potentials were applied: one to the dihedral potential and the other is applied to the total potential (Wang et al., 2011). In our simulation system, we used the following aMD parameters:

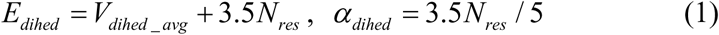

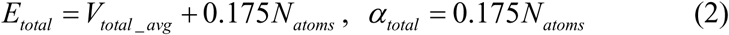

where the *E_dihed_* and the *E_total_* are the thresholds of the dihedral and total potential in the aMD, respectively; *V_dihed_avg_* and *V_total_avg_* are the dihedral and potential energy calculated from the last 10 ns of the cMD, respectively; the *N_res_* and the *N_atom_* are the protein residue numbers and the atom number of the whole system, respectively. These parameters were suggested by previous publications for proper acceleration of conformational changes in global proteins (Miao et al., 2014; Pierce et al., 2012).

## QUALIFICATION AND STATISTICAL ANALYSIS

RMSD, RMSF and interaction energy were calculated with corresponding VMD plugins. The clustering analysis in Figure 3 was done with Carma program. The DCCM in Figure 4C was calculated with the following formulation:

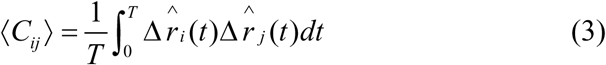

where ⟨*C_ij_⟩* is the average cross-correlation *of* the *i^th^* and *j^th^* atoms during the length of time (*T*) over which we calculated the matrix, and 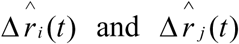 are the displacement of the *i^th^* and *j^th^* atoms at time t.

The free energy landscape of the peptide in the aMD was calculated by two-dimension histogram of the two chosen reaction coordinates, *ξ1* and *ξ2:*

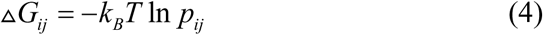

where Δ*G_ij_* is the free energy of the bin*_ij_* in the histogram which is binned into a *i*×*j* matrix according to the two reaction coordinates, *p_ij_* is the number of the snapshots falling into the bin*_ij_* (the probability distribution of the reaction coordinates), *k_B_* is the Boltzmann constant, and *T* is the simulation temperature.

In order to obtain the reweighted canonical free energy surface from an aMD, the probability distribution *p_ij_* calculated by two-dimension histogram in the aMD is reweighted to canonical ensemble distribution by:

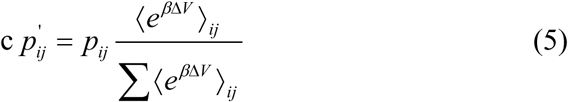

where 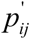 is the reweighted probability distribution in the *bin_ij_*, ⟨*e^βΔV^*⟩_*ij*_ is the ensemble-averaged Boltzmann factor of ΔV for simulation frames falling into the *bin_ij_*. Σ⟨*e^βΔV^*⟩_*ij*_ is the sum of the ⟨*e^βΔV^*⟩_*ij*_ in all bins of the two-dimension histogram. Equation 4 provides an exponent average algothrim for reweighting. In the present study, we use Maclaurin series algorithm to approximate the exponential term as the following:

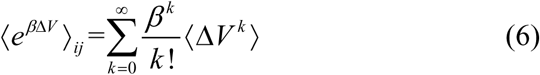

Five to ten order (k = 5∼10) expansion were used here to obtain the best reweighting result. After the reweighted probability *p_ij_* is calculated, the reweighted free energy surface is calculated by:

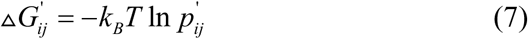

## REFERENCES

Achour, A., Michaelsson, J., Harris, R.A., Odeberg, J., Grufman, P., Sandberg, J.K., Levitsky, V., Karre, K., Sandalova, T., and Schneider, G. (2002). A structural basis for LCMV immune evasion: subversion of H-2D(b) and H-2K(b) presentation of gp33 revealed by comparative crystal structure. Analyses. Immunity 17, 757–768.

Bjorkman, P.J., Saper, M.A., Samraoui, B., Bennett, W.S., Strominger, J.L., and Wiley, D.C. (1987a). The Foreign Antigen-Binding Site and T-Cell Recognition Regions of Class-I Histocompatibility Antigens. Nature 329, 512–518.

Bjorkman, P.J., Saper, M.A., Samraoui, B., Bennett, W.S., Strominger, J.L., and Wiley, D.C. (1987b). Structure of the Human Class-I Histocompatibility Antigen, Hla-A2. Nature 329, 506–512.

Borbulevych, O.Y., Piepenbrink, K.H., and Baker, B.M. (2011). Conformational Melding Permits a Conserved Binding Geometry in TCR Recognition of Foreign and Self Molecular Mimics. Journal of immunology 186, 2950–2958.

Borbulevych, O.Y., Piepenbrink, K.H., Gloor, B.E., Scott, D.R., Sommese, R.F., Cole, D.K., Sewell, A.K., and Baker, B.M. (2009). T Cell Receptor Cross-reactivity Directed by Antigen-Dependent Tuning of Peptide-MHC Molecular Flexibility. Immunity 31, 885–896.

Dror, R.O., Dirks, R.M., Grossman, J.P., Xu, H., and Shaw, D.E. (2012). Biomolecular simulation: a computational microscope for molecular biology. Annual review of biophysics 41, 429–452.

Foloppe, N., and MacKerell, A.D. (2000). All-atom empirical force field for nucleic acids: I. Parameter optimization based on small molecule and condensed phase macromolecular target data. Journal of computational chemistry 21, 86–104.

Glithero, A., Tormo, J., Doering, K., Kojima, M., Jones, E.Y., and Elliott, T. (2006). The crystal structure of H-2D(b) complexed with a partial peptide epitope suggests a major histocompatibility complex class I assembly intermediate. The Journal of biological chemistry 281, 12699–12704.

Gras, S., Burrows, S.R., Turner, S.J., Sewell, A.K., McCluskey, J., and Rossjohn, J. (2012). A structural voyage toward an understanding of the MHC-I-restricted immune response: lessons learned and much to be learned. Immunol Rev 250, 61–81.

Hamelberg, D., Mongan, J., and McCammon, J.A. (2004). Accelerated molecular dynamics: a promising and efficient simulation method for biomolecules. The Journal of chemical physics 120, 11919–11929.

Insaidoo, F.K., Borbulevych, O.Y., Hossain, M., Santhanagopolan, S.M., Baxter, T.K., and Baker, B.M. (2011). Loss of T cell antigen recognition arising from changes in peptide and major histocompatibility complex protein flexibility: implications for vaccine design. The Journal of biological chemistry 286, 40163–40173.

Kass, I., Buckle, A.M., and Borg, N.A. (2014). Understanding the structural dynamics of TCR-pMHC complex interactions. Trends Immunol 35, 604–612.

Knapp, B., Demharter, S., Esmaielbeiki, R., and Deane, C.M. (2015). Current status and future challenges in T-cell receptor/peptide/MHC molecular dynamics simulations. Briefings in bioinformatics.

Mazza, C., and Malissen, B. (2007). What guides MHC-restricted TCR recognition? Semin Immunol 19, 225–235.

Miao, Y., Sinko, W., Pierce, L., Bucher, D., Walker, R.C., and McCammon, J.A. (2014). Improved Reweighting of Accelerated Molecular Dynamics Simulations for Free Energy Calculation. J Chem Theory Comput 10, 2677–2689.

Morikis, D., and Lambris, J.D. (2004). Physical methods for structure, dynamics and binding in immunological research. Trends Immunol 25, 700–707.

Narzi, D., Becker, C.M., Fiorillo, M.T., Uchanska-Ziegler, B., Ziegler, A., and Bockmann, R.A. (2012). Dynamical characterization of two differentially disease associated MHC class I proteins in complex with viral and self-peptides. J Mol Biol 415, 429–442.

Omasits, U., Knapp, B., Neumann, M., Steinhauser, O., Stockinger, H., Kobler, R., and Schreiner, W. (2008). Analysis of key parameters for molecular dynamics of pMHC molecules. Mol Simulat 34, 781–793.

Pettersen, E.F., Goddard, T.D., Huang, C.C., Couch, G.S., Greenblatt, D.M., Meng, E.C., and Ferrin, T.E. (2004). UCSF chimera - A visualization system for exploratory research and analysis. Journal of computational chemistry 25, 1605–1612.

Phillips, J.C., Braun, R., Wang, W., Gumbart, J., Tajkhorshid, E., Villa, E., Chipot, C., Skeel, R.D., Kale, L., and Schulten, K. (2005). Scalable molecular dynamics with NAMD. Journal of computational chemistry 26, 1781–1802.

Pierce, L.C., Salomon-Ferrer, R., Augusto, F.d.O.C., McCammon, J.A., and Walker, R.C. (2012). Routine Access to Millisecond Time Scale Events with Accelerated Molecular Dynamics. J Chem Theory Comput 8, 2997–3002.

Reboul, C.F., Meyer, G.R., Porebski, B.T., Borg, N.A., and Buckle, A.M. (2012). Epitope flexibility and dynamic footprint revealed by molecular dynamics of a pMHC-TCR complex. PLoS computational biology 8, e1002404.

Rudolph, M.G., Stanfield, R.L., and Wilson, I.A. (2006). How TCRs bind MHCs, peptides, and coreceptors. Annu Rev Immunol 24, 419–466.

Saini, S.K., Ostermeir, K., Ramnarayan, V.R., Schuster, H., Zacharias, M., and Springer, S. (2013). Dipeptides promote folding and peptide binding of MHC class I molecules. Proceedings of the National Academy of Sciences of the United States of America 110, 15383–15388.

Scott-Browne, J.P., Crawford, F., Young, M.H., Kappler, J.W., Marrack, P., and Gapin, L. (2011). Evolutionarily conserved features contribute to alphabeta T cell receptor specificity. Immunity 35, 526–535.

Shaw, D.E., Maragakis, P., Lindorff-Larsen, K., Piana, S., Dror, R.O., Eastwood, M.P., Bank, J.A., Jumper, J.M., Salmon, J.K., Shan, Y.B., et al. (2010). Atomic-Level Characterization of the Structural Dynamics of Proteins. Science 330, 341–346.

Singharoy, A., Polavarapu, A., Joshi, H., Baik, M.H., and Ortoleva, P. (2013). Epitope Fluctuations in the Human Papillomavirus Are Under Dynamic Allosteric Control: A Computational Evaluation of a New Vaccine Design Strategy. J Am Chem Soc 135, 18458–18468.

Smock, R.G., and Gierasch, L.M. (2009). Sending Signals Dynamically. Science 324, 198–203.

Song, I., Gil, A., Mishra, R., Ghersi, D., Selin, L.K., and Stern, L.J. (2017). Broad TCR repertoire and diverse structural solutions for recognition of an immunodominant CD8+ T cell epitope. Nat Struct Mol Biol 24, 395–406.

Speir, J.A., Stevens, J., Joly, E., Butcher, G.W., and Wilson, I.A. (2001). Two different, highly exposed, bulged structures for an unusually long peptide bound to rat MHC class I RT1-Aa. Immunity 14, 81–92.

Sun, Y.P., and Liu, Q. (2015). Differential structural dynamics and antigenicity of two similar influenza H5N1 virus HA-specific HLA-A*0201-restricted CLT epitopes. Rsc Adv 5, 2318–2327.

Turner, S.J., Doherty, P.C., McCluskey, J., and Rossjohn, J. (2006). Structural determinants of T-cell receptor bias in immunity. Nature Reviews Immunology 6, 883–894.

Turner, S.J., Kedzierska, K., Komodromou, H., La Gruta, N.L., Dunstone, M.A., Webb, A.I., Webby, R., Walden, H., Xie, W., McCluskey, J., et al. (2005). Lack of prominent peptide-major histocompatibility complex features limits repertoire diversity in virus-specific CD8+ T cell populations. Nat Immunol 6, 382–389.

Tynan, F.E., Borg, N.A., Miles, J.J., Beddoe, T., El-Hassen, D., Silins, S.L., van Zuylen, W.J.M., Purcell, A.W., Kjer-Nielsen, L., McCluskey, J., et al. (2005a). High resolution structures of highly bulged viral Epitopes bound to major histocompatibility complex class I - Implications for T-cell receptor engagement and T-cell immunodominance. Journal of Biological Chemistry 280, 23900–23909.

Tynan, F.E., Burrows, S.R., Buckle, A.M., Clements, C.S., Borg, N.A., Miles, J.J., Beddoe, T., Whisstock, J.C., Wilce, M.C., Silins, S.L., et al. (2005b). T cell receptor recognition of a 'super-bulged' major histocompatibility complex class I-bound peptide. Nat Immunol 6, 1114–1122.

Valkenburg, S.A., Josephs, T.M., Clemens, E.B., Grant, E.J., Nguyen, T.H., Wang, G.C., Price, D.A., Miller, A., Tong, S.Y., Thomas, P.G., et al. (2016). Molecular basis for universal HLA-A*0201-restricted CD8+ T-cell immunity against influenza viruses. Proceedings of the National Academy of Sciences of the United States of America 113, 4440–4445.

Velloso, L.M., Michaelsson, J., Ljunggren, H.G., Schneider, G., and Achour, A. (2004). Determination of structural principles underlying three different modes of lymphocytic choriomeningitis virus escape from CTL recognition. Journal of immunology 172, 5504–5511.

Wan, S., Coveney, P., and Flower, D.R. (2004). Large-scale molecular dynamics simulations of HLA-A*0201 complexed with a tumor-specific antigenic peptide: can the alpha3 and beta2m domains be neglected? Journal of computational chemistry 25, 1803–1813.

Wang, Y., Harrison, C.B., Schulten, K., and McCammon, J.A. (2011). Implementation of Accelerated Molecular Dynamics in NAMD. Computational science & discovery 4.

